# Refactoring the Genetic Code for Increased Evolvability

**DOI:** 10.1101/128058

**Authors:** Gur Pines, James D. Winkler, Assaf Pines, Ryan T. Gill

## Abstract

The standard genetic code is robust to mutations and base-pairing errors during transcription and translation. Point mutations are most likely to be synonymous or preserve the chemical properties of the original amino acid. Saturation mutagenesis experiments suggest that in some cases the best performing mutant requires a replacement of more than a single nucleotide within a codon. These replacements are essentially inaccessible to common error-based laboratory engineering techniques that alter single nucleotide per mutation event, due to the extreme rarity of adjacent mutations. In this theoretical study, we suggest a radical reordering of the genetic code that maximizes the mutagenic potential of single nucleotide replacements. We explore several possible genetic codes that allow a greater degree of accessibility to the mutational landscape and may result in a hyper-evolvable organism serving as an ideal platform for directed evolution experiments. We then conclude by evaluating potential applications for recoded organisms within the synthetic biology field.

**Significance Statement:** The conservative nature of the genetic code prevents bioengineers from efficiently accessing the full mutational landscape of a gene using common error-prone methods. Here we present two computational approaches to generate alternative genetic codes with increased accessibility. These new codes allow mutational transition to a larger pool of amino acids and with a greater degree of chemical differences, using a single nucleotide replacement within the codon, thus increasing evolvability both at the single gene and at the genome levels. Given the widespread use of these techniques for strain and protein improvement along with more fundamental evolutionary biology questions, the use of recoded organisms that maximize evolvability should significantly improve the efficiency of directed evolution, library generation and fitness maximization.

## Introduction

The deciphering of the genetic code has yielded many insights into its organization and evolution (1). Notably, the standard genetic code was found to be robust to many single nucleotide replacements (SNRs) and that similar codons code for amino acids with related properties (Fig. 1A) (2–5). Theories around the origin of the standard code organization focus on three main ideas: The first is the stereochemical theory, claiming that the current genetic code was formed by primordial interactions between codons (or anticodons) and amino acids (6). The second, termed the coevolution theory, points to the fact that amino acids sharing biosynthetic pathways are generally assigned to similar codons and suggests that the genetic code today reflects this coevolution (7–9). The third theory hypothesizes that the genetic code was under selection to minimize deleterious changes in physicochemical properties caused by mutations and mistranslations (3, 10). A combination of all three theories is also a possibility (11). Regardless of the exact mechanism, the modern standard genetic code is indeed relatively resistant to SNRs and buffers their effect (2, 12–14). As a result, a given amino acid can only be converted on average into 6.1 others (including the stop codon) without multiple nucleotide changes. While this property is favorable for buffering mutational effects in free-living organisms, it limits the effectiveness of mutagenesis techniques relying on single nucleotide substitutions to alter coding sequences.

**Figure 1.**
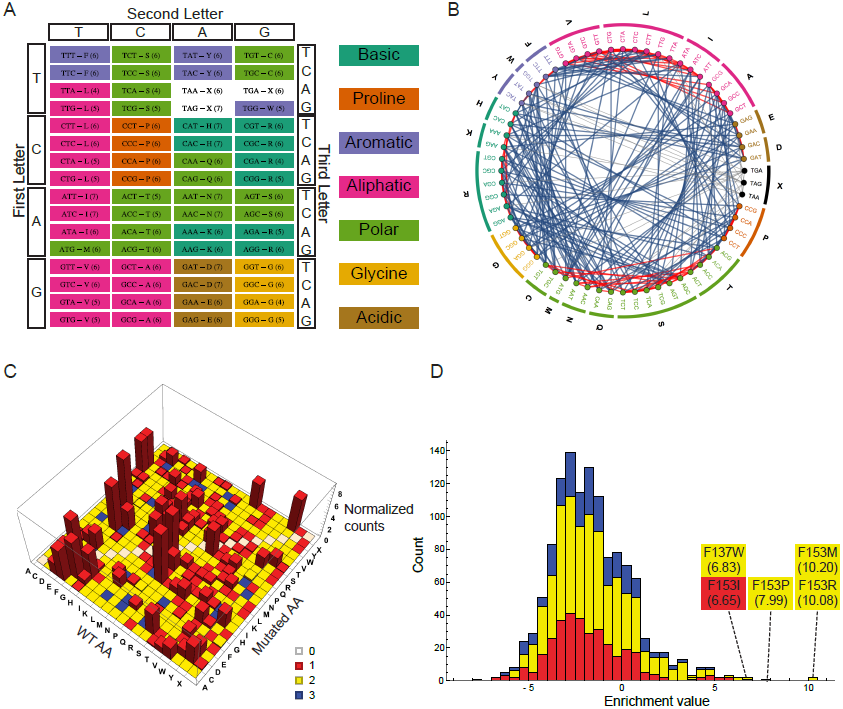
The properties of the standard genetic code. (A) The standard genetic code. Colors indicate the amino acid chemical classes as depicted on the right site of the chart. Stop codons are colored white and are denoted by “X”. Numbers in parentheses indicate the number of SNR-accessible unique amino acids for each codon. (B) The codon SNR accessibility plot of the standard code. All 64 codons are grouped according to their corresponding amino acids which in turn are clustered by their chemical classes (colors are as in A). Edges connecting two codons indicate that these two codons are within an SNR distance. Edge color corresponds to whether the amino acid change is within the same chemical class (red) or between classes (blue). Edges connecting to stop codons are shaded in gray. (C) Analysis of previously reported resistance mutations. The X and Y axes correspond to the wild-type and mutated amino acids in resistance-conferring genes. Colors indicate the minimal number of nucleotide changes needed within the corresponding codons. The Z axis represents normalized counts of the mutations (adapted from Winkler et al. 2016 (43)). Colors indicate the minimal number of nucleotide replacements needed for the transition from WT amino acid to the mutated one. Note that in the vast majority of mutations may be explained by a single nucleotide replacement within a codon. (D) A stacked view histogram of scanning saturation mutagenesis data from Garst et. al. (32), showing the mutational fitness landscape following the incubation of a *folA* library with trimethoprim. Colors are as in C. Note that the four most enriched mutants require two nucleotide replacements within the codon (enrichment values and mutation identity are indicated).

Saturation mutagenesis techniques provide full access to all amino acids by using degenerate codons covering a comprehensive collection of amino acids (15–17). Such approaches frequently identify multiple nucleotide replacements (MNRs) within a codon to be superior to SNRs in a variety of contexts (16, 18–26). Recent technological advances in DNA synthesis and sequencing have enabled more detailed surveys of a complete systematic saturation mutagenesis of a gene, permitting the interrogation of the full mutational landscape at single amino acid resolution (27–29). Data from such experiments are still relatively rare but support the notion that the highest peaks in the mutational landscapes can require MNRs that are inaccessible in reasonable time frames using SNRs only (30–32).

For investigations of the combinatorial space, saturation mutagenesis may be performed iteratively, but requires *a priori* knowledge of sites of interest (33–35). Computational approaches intelligently select sites of interest, thus reducing the need to survey all possible mutations (36–38), but the risk of missing important sites still exists; the desired phenotype may consist of multiple unpredictable changes in genes and other genetic elements. As a result, adaptive laboratory evolution experiments are still the gold standard for identifying desired phenotypes when the underlying genotype-phenotype relationship is unclear. However, laboratory evolution and error-prone PCR, commonly used for protein engineering, cannot access the full amino acid repertoire since they are primarily limited to SNRs to alter protein coding sequences, and thus might not be able to access the genotypes at the global fitness maximum. While further improvement is feasible with further secondary mutations, the multiple rounds of diversity generation followed by selection slows the process of strain or protein improvement significantly (39–42).

Here we suggest several reorderings of the genetic code that will enable significantly more comprehensive and rapid exploration of fitness landscapes and simplify the identification of beneficial mutations. We apply two computational approaches for the generation of such genetic codes and discuss a range of alternative codes that have altered SNR accessibilities. Subsequently, we recode the set of essential *E. coli* genes using these generated codes to evaluate the high-level impact of genetic code refactoring. We then discuss both the technological improvements required for implementing these hypothetical codes in a living organism and potential biotechnological applications for novel, highly evolvable genetic codes.

## Results

### The amino acid accessibility

At the amino acid level, the standard genetic code allows an SNR to convert a codon into alternative codons encoding 6.1 unique amino acids. The distribution is not wide, as both the standard deviation (0.69) and range (5 to 7 amino acids) are low as a result of the constraints placed on mutational accessibility due to the architecture of the tri-nucleotide genetic code. On average, 5.53 different chemical classes are accessible by the same mutations (See Methods for amino acid classification source).

Figure 1A depicts the standard genetic code organization with the amino acids colored according to their chemical classes, and numbers in parentheses indicate the number of unique SNR-accessible amino acids. An additional manner of illustrating the SNR accessible amino acids is shown in Fig. 1B. Here, the codons are ordered in a circle, grouped according to the encoded amino acids and classes. Edges connecting two codons represent SNR accessibilities, and edge color indicates whether amino acid conversion is within (red) or between (blue) chemical classes. This plot highlights the notion that many SNR-accessible amino acids are chemically similar as depicted by the multitude of red edges. Edge distribution of all genetic codes presented in this study are shown in Fig S2.

To investigate the potential effect of mutation buffering that favors SNR-induced amino acid replacements, we analyzed a recently published collection of 2679 amino acid changes generated by various random methods associated with diverse resistance phenotypes (43). The results show that almost all mutations found could be explained by a single nucleotide replacement, with double- and triple nucleotide replacements being required for at most 3.5% of all detected mutations (Fig. 1C). The number of nucleotide replacements required for inter-amino acid conversion and their observed frequency in the data is strongly anti-correlated (P < 10^−10^, Spearman), comporting well with previous findings (44). Taking into account that this database is related to mutational response to stress, known to be associated with elevated mutation rate, increases the significance of these findings ever further (45–47). These findings highlight the confined space evolution is allowed to explore since it is mostly limited to SNRs.

Recently, we reported a novel method for high throughput, genome-wide, single amino acid level, genome editing in *E. coli* (32). A complete scanning saturation mutagenesis of the DHFR gene, *folA*, and incubation with the DHFR-specific inhibitor trimethoprim, was performed to find resistant mutants. Multiple mutants were found to be enriched following trimethoprim treatment, including several sites that were previously reported (39, 48). However, many of those sites, including the mutants with the highest fitness values (Fig. 1D), had novel MNR mutations, since most experiments were previously done by SNR-based methods such as directed evolution (39, 48). These results are in line with other findings, suggesting that MNRs are more effective at phenotype improvement than SNRs found by random mutagenesis methods (16). Other saturation mutagenesis libraries also show MNR superiority over SNRs under several adaptive conditions (30–32, 44). Taken together, these examples along with the fundamentally conservative nature of the genetic code support the hypothesis that sometimes a radical change in the amino acid characteristics is required for a drastic shift in the corresponding protein’s properties and that such changes are simply not available using SNRs alone.

### Generating alternative genetic codes

Having identified a critical factor limiting the effectiveness of current directed evolution tools, uncovering possible genetic changes that can reduce the buffering capacity of the standard genetic code is of interest. So far, *E. coli* has been engineered to lack the TAG stop codon, and a larger scale ongoing effort to remove seven codons from the genome has been reported (49, 50). These efforts establish the conceptual feasibility of altering the code and producing a viable organism with a more “evolvable” code if it can be designed and implemented successfully. One approach to generating new candidate codes is through the use of a genetic algorithm that evolves new codes according to user-defined criteria. Genetic algorithms have been used to optimize other properties of genetic codes in the past, typically focusing on robustness (51) or propagation of error to the protein level (52). The principal requirement for implementing a genetic algorithm is defining a fitness function F that can be maximized to select amongst a population of randomly modified codes to identify those encoding our desired characteristics outlined above.

The first component F_{unique} of our proposed fitness function is simply the average cardinality of the set of amino acids accessible by each codon within the genetic code using SNRs (Equation 1), where num(*) represents the cardinality operator, AA(C) represents the amino acid encoded by a codon C, D(X,Y) a hamming distance function, and N the number of codons in the genetic code. The second component F_{ratio} (Equation 2) is calculated by first determining the number of times an SNR will convert any other amino acid into a given amino acid, and then dividing the minimum and maximum value to obtain the ratio of interest for a code with M total amino acids. Codes with more even distributions of codons between amino acids will have F_{ratio} values closer to one. The final component of F is simply the number of non-distinct chemical classes accessible by SNRs, denoted as F_{chem} (Equation 3). The final fitness function (Equation 4) is then taken as the product of these components normalized by their theoretical maximums to avoid inadvertently favoring improvement in a specific area over the others (see Methods).

However, one limitation of this approach is that there is no guarantee that the final ensemble of improved codes has reached the global fitness optimum, given the stochastic nature of the algorithm and extremely large number of possible valid codes. Calculation of F(C) is also O(N^2^) in nature because accessibility between all pairs of amino acids must be computed, and becomes computationally limiting if the number of candidate codes per round of selection is large.

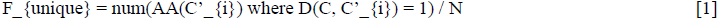

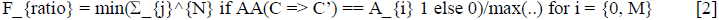

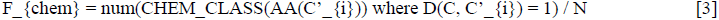

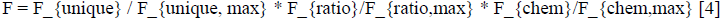

Applying the selection procedure outlined here (see Methods) yields genetic codes with a significantly higher level of codon accessibility both concerning the number (Fig. S1A) and the chemical diversity of the SNR-resulting amino acids (optimized (OPT) code, Fig. 2A, note the decrease in red edges). The recoded genetic code includes a single stop codon manually added after the selection procedure. Increased amino acid accessibility naturally leads to a concomitant decrease in robustness to mutation by allowing SNRs to lead to a wider variety of amino acid substitutions (Fig. S1A), given the inverse relationship of these properties. The bias in codon distribution in the standard code between different amino acids had been flattened such that the number of codons assigned for amino acids is between 2 to 4, compared to from 1 to 6 in the standard code. Adding a native codon restriction to ensure that each amino acid has at least one native codon produced the OPT-NR code, which is slightly less optimized but still with a higher degree of fitness than the standard code (as defined by the genetic algorithm, Fig. 2B, S1B). The distribution of the inter- and intra-chemical class SNR conversions is shown in figure S2.

**Figure 2.**
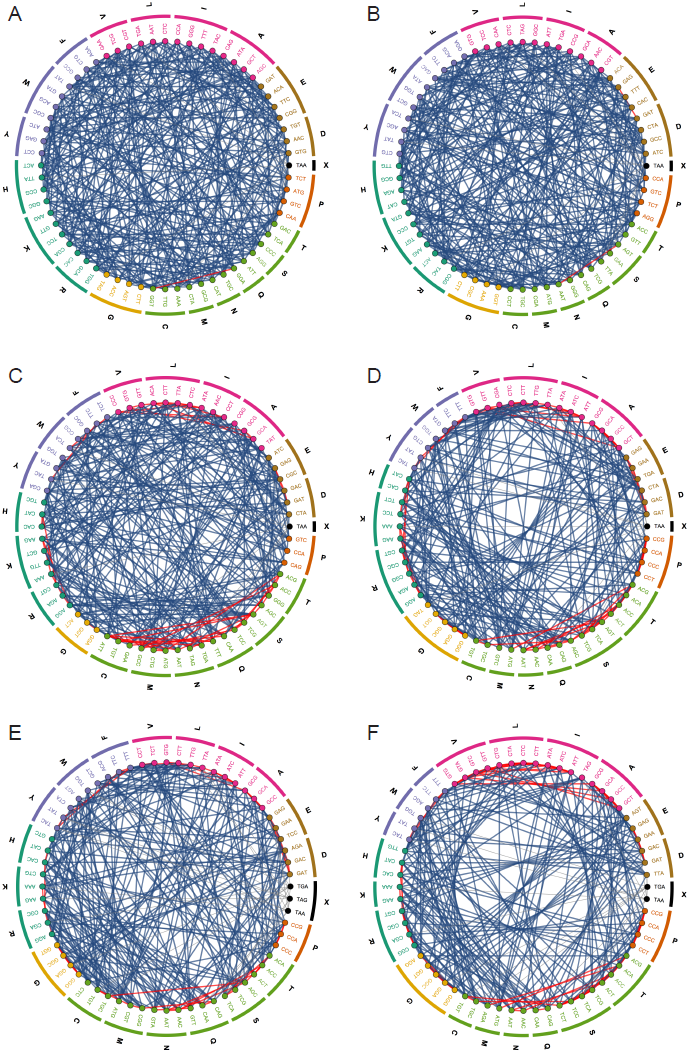
Accessibility plots of the genetic codes presented in this study. Plots are organized as in Fig. 1B. (A) the optimal code (OPT). (B) the optimal code with the imposed native codon restriction (OPR-NR). (C) change minimizing code utilizing a linear penalty for changes (CMC). (D) change minimizing code utilizing a power of 2 penalty for changes (CMC^2^). (E) genetic code derived from a recursive approach of amino acid reassignment (REC). (F) the resulting genetic code of reassigning the seven free codons as described by Ostrov et. al. (50)

While these genetic codes represent maximal accessibility via SNRs, they require 60 and 43 codon reassignments for the OPT and OPT-NR, respectively. Genetic code refactoring on such a scale is not likely to be feasible in the near future due to the complexities of simultaneously recoding the genes while preserving key properties such as secondary structure, ribosome binding sites, and other features (see Discussion). A second fitness function that penalizes codon reassignments in the effort to balance maximal accessibility with a minimal number of reassignments resulted in another two genetic codes termed CMC (change minimizing code) and CMC^2^. These codes require 30 and 9 reassignments using linear and power penalties but with 25.1% and 10.2% improved SNR accessibility (F_{unique}) relative to the standard code nonetheless (Fig. 2C and D). Genetic code tables and the chemical difference of SNR-accessible amino acids are shown in Figures S1C, S1D, and S2. Given that minimal changes substantially improve each component of our fitness function (Fig. S3), it may be possible to create a platform organism with a small number of reassigned codons that achieves the lion’s share of possible accessibility improvement.

A second approach for increased evolvability by modifying the genetic code, one that might prove more practical, is a stepwise reassignment of amino acids. This process entails the identification of the optimal amino acid replacement(s) and then re-evaluating the resulting altered code until no further single-step improvement is possible. Since there are often many amino acid reassignments that yield equivalent fitness improvements, we implemented a recursive branching approach to exhaustively evaluate all potential single-step reassignment procedures. Using this method, the code’s fitness reached its maximum after 16 recursive rounds and resulted in a 14.8% improvement in unique amino acid accessibility compared to the standard code (Fig. 2E, S1E, S3A). This code exhausted all available single-step improvements, demonstrating that multiple simultaneous mutations are necessary to improve the code further. While this approach resulted in less evolvable codes than the OPT codes (see below), this approach may be more practical for near future experimental evaluations by enabling iterative codon reassignment rather than simultaneous wholesale code engineering.

### Maximizing Accessibility for Existing Recoded Organisms

Genome-wide multiple reassignments are not trivial and may result in a non-viable organism that is challenging to test and modify. There are currently only a handful of organisms with artificially modified genetic codes and the attendant genome refactoring, one of which may be ideal to test our approach for increasing SNR accessibility. Ostrov *et. al*. recently reported on the ongoing effort to engineer a 57 codon E. coli genome (50), with the seven codons replaced with their synonymous counterparts. We applied our genetic algorithm to reassign these seven codons for fitness maximization (Fig. 2F, S1F) and generated a code with a 7.7% increase in unique amino acid accessibility and 9.6% increase in chemical class diversity obtainable by SNRs. While these differences are small, they represent a significant increase in the complexity accessible by random mutation and allow for new paths on the fitness landscape of interest to be explored. The tRNA reassignments required for the proposed code should not affect the physiology of the final rE. coli-57 strain since these codons are deleted from this genome. Following successful reassignment, codons may be gradually reintroduced to the genome employing MAGE or other methods (49, 53, 54). This process and the attendant debugging required will enable the understanding of the reassignment design rules that may finally lead the synthesis of a rationally re-designed genome employing a novel genetic code.

### Genetic Code Analysis

The genetic codes described here can be compared both at the levels of code fitness and by the broader term of genome evolvability. Figure 3A places all the genetic codes outlined in this study in a 3-dimensional space composed of the three parameters that were subject to optimization, namely number of unique SNR-accessible amino acids, their chemical diversity and the distribution of the number of codons assigned for an amino acid. Every genetic code is represented as a bubble with its size proportional to the number of the required reassignments. The OPT, OPT-NR and CMC codes are partially overlapping and clustered close to the most optimized corner of the space. The recursive code is less optimized but is still improved relatively to the CMC code that utilizes a power penalty for reassignments. The recoded rE. coli-57 code represents a different combination of optimizations than its neighbor, the CMC^2^ code and requires the lowest number of reassignments. 2-dimensional versions of figure 3A are depicted in Figures S4A-C.

**Figure 3.**
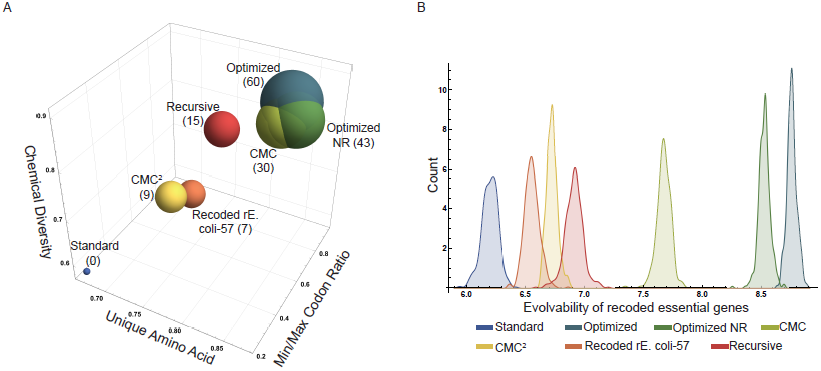
Comparisons of the genetic codes presented in this study. (A) A 3-dimensional bubble chart with the axes corresponding to the three parameters that were selected for optimization. Bubble size corresponds to the number of codon-amino acid reassignments required for each genetic code, which is also indicated in parentheses. (B) Genome evolvability of the presented codes as computed by the accessibility of the recoded collection of the E. coli essential genes.

As the genetic code and content are inextricably intertwined in living organisms, the distribution of codons observed in living organisms is the result of continual selection over billions of years for genetic stability, efficient transcription, translation, and regulation. Recoding genetic material using these artificial codes, therefore, must preserve inherent structural and sequence features found in the genome as much as possible. To evaluate the genome-wide effects of recoding, we implemented a resequencing algorithm that seeks to minimize predicted changes in folding energy and secondary structure (see Methods) and then recoded the set of known essential E. coli genes using a range of generated genetic codes for analysis. We therefore defined an “evolvability” score calculated according to Equation 5, which iterates over the length of a protein and sums the number of unique amino acids that are accessible from a given codon i (D = 1, required for an SNR to convert C to C’) and normalizes the result by the length of the protein P.

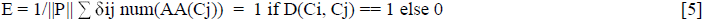

The wild-type genes were found to have lower, more variable evolvability scores compared to the genes recoded using the generated codes in rough concordance to their calculated fitness scores (Fig. 3B, S3A); higher fitness codes permit greater sequence flexibility compared to the more constrained standard code. However, since synonymous codon recoding is now feasible on a large scale (49, 53, 55), it may be possible to selectively recode portions of the genome such that they use these reassigned codons more frequently.

## Discussion

The modern genetic code buffer errors both in terms of mutations and translation and as a result it limits SNR-based amino acid accessibility (Fig. 1A, B) (12–14, 56). This inherent property of the genetic code significantly limits error-based engineering methods such as adaptive laboratory evolution and error-prone PCR, preventing the exploration of many adaptive mutations, including at times the discovery of the most favorable fittest mutant (Fig. 1C, D). Here we describe more flexible genetic codes with access to larger portions of the full fitness landscape. While the codes discovered by our genetic algorithm presented a high degree of accessibility, they required multiple amino acid reassignments (Fig. 2, 3, S1). Using a second approach, we adopted a stepwise strategy with each step re-assigning an amino acid-codon pair in a manner that increases accessibility the most. This resulted in a less accessible final genetic code compared to the optimized ones discovered using the genetic algorithm, but has the advantage of the gradual modifications and might prove to be more practical experimentally (Fig. 2E, 3, S1E). Finally, we reassigned the seven free codons of the recently reported rE.coli-57 (50), which increased unique amino acid accessibility by 7.7% (Fig. 2F, 3, S1F) and is currently the most practical approach to test the hypothesis described here. Altogether, the genetic codes presented here demonstrate a significantly higher level of fitness regarding the number of unique SNR-accessible amino acids, their chemical properties, and the codon assignment distribution (Fig. 3A). This fitness increase may be further translated to the consequent increase in the overall evolvability at the genome level when essential genes are recoded according to these genetic codes (Fig. 3B). In general, the marginal value of codon reassignments, especially for improvements in chemical class accessibility, decreases rapidly when using the genetic algorithm (Fig. S3B), suggesting that the optimal effort-reward tradeoff must be determined to the extent of re-engineering that can be tolerated.

The physical engineering of a recoded organism will require a genome-wide recoding and reassignment of most, if not all tRNAs. A successful construction is depending mainly on the following three topics: Genome synthesis, genome recoding and the engineering of the translational mechanism. The cost of DNA synthesis is continuously decreasing, enabling large scale assemblies to be executed in laboratory environments (57, 58). Whole genome chemical synthesis was previously shown on *Mycoplasma mycoides* (59), and recently a smaller version of this genome was shown to be viable, removing many of its nonessential genes (60). Moreover, genome synthesis has been successfully demonstrated in a variety of organisms spanning from synthetic viral genomes to yeast chromosomes and mouse mitochondria (61–63). Notably, leading members of the synthetic biology community have recently proposed to synthesize the human genome, following the “learn through building” paradigm (64). Genetic code engineering and codon reassignment were shown before, mainly for genetic code expansion and incorporation of nonnatural amino acids (49, 65–68). These experiments focused on reassigning one or few codons at a time, but they demonstrate the feasibility of such feats. For a recoded genome involving multiple tRNA reassignments, multiple orthogonal tRNAs need to be identified and non-straightforward additional engineering of associated proteins such as aminoacyl transferases are required. Even the successful synthesis of the first step in our recursive approach is currently challenging. Engineering constraints such as these can be accounted for by imposing penalizing or eliminating artificial codes that require complex underlying interventions to implement.

The viability of an entirely recoded organism is not assured; the challenge of bootstrapping an altered code and debugging the resulting deleterious or lethal mutations required remains an unsolved challenge. We still are not familiar with the complete functionalities of the full genomic sequence of an organism and a complete recoding may result in a severely impaired fitness. Though it might be feasible that once a recoded organism is constructed, adaptive evolution can be used to select for genome remodeling that improves strain fitness closer to wild-type levels. Recoding a whole genome with amino acid reassignments will probably prove to be significantly more complicated than previously reported synonymous reassignments (49, 50). Finally, the genetic codes proposed here eliminate the robustness enjoyed by the standard code and disrupt the fine balance between robustness and evolvability (69, 70). This may lead to a disruption of the genomic integrity that decreases host viability. However, given that the estimates of *E. coli*’s point mutation rate are generally on the order of 10^−10^/bp-generation (71), this effect will be more obvious under stress-induced mutagenesis or exposure to DNA mutagens than replication in laboratory timescales.

Any organisms using an artificial coding scheme for DNA-protein translation have the implicit benefit of strong bio-control. Environmental releases of an organism altered in this fashion cannot easily express environmental DNA and cannot transfer heterologous DNA to other microbes due to the drastic differences in their coding schemes. On the other hand, considering its resistance to viral infections (68), a release of such organism to the environment is a concern. Suitable safety measures such as dependence on exogenously-supplied ligands, suicide circuits or nuclease-based DNA destruction should be implemented (72, 73). Another approach is to induce addiction to an environmentally unavailable nonstandard amino acid, ensuring that its escape from the laboratory environment would result in cell death (74, 75). This seems particularly fitting to the case presented here since it requires genome recoding and freeing at least a single codon for reassignment purposes, which could be easily accommodated into such designs.

Here we present the concept of refactoring the genetic code with the aim of increasing evolvability by optimizing the SNR-accessible amino acids regarding both the number of unique amino acids and their chemical properties. We added a third parameter to our fitness function to avoid codon bias (Fig. 3A). It is possible to alter these settings, such as using different chemical property classes or adding more dimensions such as amino acid size, secondary structure propensity, molecular weight, etc. Other options include directing the genetic code for having a different bias than the one that exists in the standard code or making a less evolvable genetic code, for maintaining desirable traits.

A successful synthesis of such an organism may result in extreme evolvability with increased accessibility to a significantly larger portion of the fitness landscape, enhancing the effect of SNR-dependent directed evolution methods for protein engineering and strain design purposes. Once the desired phenotype is isolated, the corresponding amino acid sequence can be reverse-engineered to the standard code for incorporation in natural, more stable organisms. In addition, such an organism may serve as a platform for studying more fundamental questions such as the evolvability of the genetic code and evolutionary robustness.

## Methods

### Computational Details

All simulations were run on a T430 Thinkpad (Windows 8.1, 16 GB RAM) in Python 2.7.11. The genetic algorithm described below was run using the PyPy 5.4.1 interpreter, while all other scripts were run using the standard Python 2.7.11 interpreter. RNAFold (76) was used to compute secondary structures and folding energies for all sequences. Visualizations were generated using Mathematica 10. Code and data used in this study are deposited in the https://bitbucket.org/jdwinkler/genetic_code_generator/ repository for download.

### Calculation of Codon Accessibility

The average number of SNR-accessible amino acid transitions was calculated by computing the number of codons encoding different unique amino acids with a Hamming distance of 1 per amino acid, and then averaging the result. For the standard E. coli genetic code, each codon can be converted into 6.1 unique amino acids by SNRs on average.

### Genetic Algorithm for Code Generation

The genetic algorithm was used to evolve the native genetic code (referred to as the standard in the text) into codes that maximize the number of alternative amino acids with distinct chemical classes accessible by mutating a codon C with only SNRs. The full implementation of the genetic algorithm is provided in the repository above; only a general description is included here.

The fitness function used for the genetic algorithm has three components. Firstly, the number of unique amino acids accessible by SNRs averaged over all codons in the genetic code (F_{unique}) is used to select for codes where SNRs lead to the greatest diversity of amino acids that can be obtained by mutating a single codon. Next, the number of mutations leading to the least accessible amino acid divided by the number of mutations leading to the most accessible amino acid (F_{ratio}) is included as part of the fitness function is used to prevent situations where nearly all codons are assigned to a single amino acid; codes with higher values of F_{ratio} will have more even distributions of codons between amino acids. Finally, we include a chemical diversity score F_{chem} that counts the number of non-distinct amino acid chemical classes accessible from a given codon by SNRs. The overall fitness function is the product of these individual components (F=F_{unique}F_{ratio}F_{chem}) divided by their theoretical maximums (F_{unique, max} = 9, F_{ratio} = 1.0, F_{chem} = 9) to avoid inappropriate weighting during selection. Amino acid chemical classes definitions were obtained from the Sigma amino acid reference chart (http://www.sigmaaldrich.com). Codon reassignments can also be penalized by reducing F using a factor alpha^∧^N, where N = 1 for linear penalties and N = 2 for square penalties, resulting in the Change Minimizing Code (CMC) and CMC^2^, respectively.

Each simulation utilized 2500 individuals and lasted 1000 rounds. For codes that exclude the stop codon from the genetic algorithm procedure, TAG and TGA were arbitrarily reassigned to asparagine and TAA was reserved to be the sole stop codon available. The codes for the next round of selection are generated using random mutation of the top 10% in a cyclical manner. The previous winner is also retained in the population undergoing selection to avoid fitness regression during the procedure. Since our goal was to select codes with optimized chemical diversity, an additional check was added to only accept new codes that had at least the same level of diversity (F_{chem}) as the previous best observed in the simulation. Codes can be subject to a range of constraints, including fixed amino acids, requiring at least one codon to retain the wild-type amino acid assignment, and permitting the removal of amino acids during the selection process. The utilization of these flags is specified in the code. During all simulations, the mutation rate was 10% per codon (i.e. 10% chance a codon would have its amino acid assignment swapped with another randomly chosen codon). Our empirical results indicated that this mutation rate generally yielded rapid improvements without trapping codes within local fitness minima due to extreme mutation rates.

### Single-step Code Generation

As an alternative to the genetic algorithm, we implemented a “best-move” based improvement scheme that iteratively improves the standard genetic code. Briefly, all possible codon reassignments for an input code are considered. To exhaustively evaluate possible best move codes, we implemented a recursive branching algorithm that generates all possible single move codes with equivalent fitness improvements until no further accessibility improvement is possible, The final ensemble of codes is then analyzed to find the set of those with maximized fitness. Stop codons are not included as part of the search but are added back after all moves have been exhausted.

### Gene Resequencing with Generated Codes

Once candidate codes were generated using the genetic algorithm or recursive approach outlined above, we next recoded the set of known *E. coli* essential genes to use these new codon-amino acid mappings. Given previous data showing that the secondary structure of the four bp of the 5’ UTR along with the first 37 bp of a coding mRNA had the largest impact on translation efficiency (77), we split the recoding process into two parts. First, the coding section of the mRNA (padded out to 39 bp) was translated according to the standard code into the corresponding polypeptide sequence; non-coding bases were not altered. The translated sequences were then converted back into a sequence of candidate codons using the generated genetic codes, yield a sequence of lists. Using the python itertools library, we built the cartesian product of these possible codon combinations for the truncated mRNA and scanned 1% of all possible assembled sequences to identify the one that best matched the wild-type folding energy and secondary structure. Ideally, this selection scheme would theoretically maintain the same transcription and translation efficiency observed for the wild-type sequence, but it does neglect the preservation of other sequence features (RBSs, for example) for simplicity. Once this initial scan is complete, areas where the recoded mRNA leader and the original mRNA sequences differ in secondary structure are identified and alternative codons from the new proposed code substituted to reduce the structural disparity.

After the recoded leader sequence is finalized, the entire candidate gene is then translated from the original standard code to the new candidate coding scheme. The process of whole gene recoding is conceptually similar to that outline for the leader sequence, but with several key differences due to the larger scale of the problem. It is not feasible to compute or scan the cartesian product of all possible codon combinations for a coding sequence due to a combinatorial explosion of candidates, so instead of analyzing the whole sequence at once, a sliding window approach is used to reduce the number of candidate sequences. The leader sequence of each essential gene is replaced with its recoded counterpart (computed as above), followed by sequence optimization of 30 bp slices at once using the same approach employed for the mRNA leader sequences. Only 0.4% of all possible candidates are screened to reduce the computational burden of sequence exploration, although it is feasible to increase the number of candidates examined depending on the availability of computational resources. Once all windows within the sequence were analyzed, there was no further adjustment to remove discrepancies between the wild-type and recoded sequence structural properties. Other more comprehensive methods are also available (50) that may produce improved recoded gene candidates.

## Acknowledgements

We thank our lab members, Andrew Garst, and Ron Milo for helpful discussions and the conceptualization of the ideas presented here. This work was funded by the U.S. Department of Energy Grant No. DE-SC008812.

